# Decrease of Functional Connectivity within the Default Mode Network by a Brief Training of Focused Attention on the Breath in Novices

**DOI:** 10.1101/2021.02.09.430388

**Authors:** Sara Trova, Yuki Tsuji, Haruka Horiuchi, Sotaro Shimada

**Affiliations:** Department of Electronics and Bioinformatics, School of Science and Technology, Meiji University, 1-1-1 Higashi -Mita, Tama-ku, Kawasaki-shi, Kanagawa 214-8571, Japan; Research and Development Initiative, Chuo University, 1-13-27, Kasuga, Bunkyo-ku, Tokyo, 112-8551, Japan

**Keywords:** electroencephalography, focused attention, default mode network, mindfulness meditation, functional connectivity

## Abstract

Experienced meditators reduce the activity of the default mode network (DMN), a brain system preferentially active when people are not engaged in specific tasks. However, the neural modulation of the DMN in novices remain largely unexplored. By using electroencephalography, we investigated the DMN functional connectivity in two groups of novices: the meditation group practiced six consecutive days of focused attention on the breath; the control group practiced only on the first and last days. After the brief training, results showed a decrease in the DMN functional connectivity between the ventromedial prefrontal cortex and the posterior cingulate cortex in theta and alpha bands during the focused attention condition, in the meditation group compared to the control group. The change in DMN functional connectivity was significantly correlated with the increase in state-level mindfulness scores. These data elucidate DMN modifications already arising at the initial stages of mindfulness meditation training in novices.

**Highlights:**

- An effect of brief meditation training on brain activity in novices was examined by using EEG.
- A six-day training of focused attention on the breath improved state-level mindfulness scores.
- Brief meditation training also reduced the functional connectivity within anterior-posterior DMN.
- The amount of change in DMN functional connectivity was significantly correlated with the subjective score.

## 1. Introduction

A growing number of mindfulness meditation studies have reported the diminished activity of the default mode network (DMN) in experienced meditators (Brewer et al., 2011; Garrison et al., 2015; Pagnoni et al., 2008). The DMN consists of multiple brain areas preferentially active when individuals are not focused on the external environment (Buckner et al., 2008; Buckner & DiNicola, 2019) and it is often associated with mind-wandering (Christoff et al., 2009; Mason et al., 2007) and self-referential processing, i.e. remembering, envisioning the future and making social inferences (Buckner et al., 2008; Buckner & Carroll, 2007). The DMN activity relies on the functional connectivity among specific cortical areas, mainly including the anterior cingulate (ACC), medial prefrontal cortex (mPFC) and posterior cingulate cortex (PCC) (Buckner et al., 2008; Greicius et al., 2003). Concurrent researches examining the neural correlates of meditation have widely associated a DMN modulation with the meditation practice. In particular, they have shown that experienced meditators relative to controls exhibit lower DMN activity during meditation (Brewer et al., 2011), and also during an active cognitive task (Garrison et al., 2015). Considering that the DMN is known to be less active during tasks requiring cognitive effort compared to rest (Buckner et al., 2008; Raichle et al., 2015), it indicates that meditation leads to reduced activity in the DMN, more than expected by general task-based deactivation. Although the aforementioned studies referred to experienced practitioners with thousands of hours of meditation experience, recent studies have also reported a modulation in DMN activity in naïve meditators after short mindfulness-based interventions (Xiao et al., 2019; Yang et al., 2019). For example, Xiao et al. (2019) reported that 8 weeks of weekly Mindfulness-Based Stress Reduction (MBSR) training (Kabat-Zinn et al., 1992) in naïve meditators reduced the activity in the posterior DMN (e.g., PCC, precuneus and cuneus). Therefore, short-term mindfulness-based interventions, similarly to long-term practitioners, are associated with reduced neural activity within specific DMN regions.

The DMN is a distributed neural network characterized by the interactions between brain regions and its activity has been often evaluated by functional connectivity (FC) among DMN-related areas. Electroencephalography (EEG) analysis, as well as magnetoencephalography (MEG), have a high temporal resolution therefore they are suitable methods for calculating functional connectivity across brain networks (Bowyer, 2016). Previous analysis revealed that the connectivity within DMN nodes at rest is ascribed to neural oscillations mainly involving alpha (8-13 Hz; Mantini et al., 2007; Tang et al., 2017), high theta (5-8 Hz) and gamma (30-80 Hz) ranges dependent on the DMN regions analyzed (Samogin et al., 2019). Similarly, EEG studies on meditation investigating the DMN-FC often reported modulations in alpha (Berkovich-Ohana et al., 2014; Fingelkurts et al., 2016) as well as in theta (Faber et al., 2004) and gamma bands (Berkovich-Ohana et al., 2014; Faber et al., 2004) in relation with the meditation practice. In a recent study using MEG (Tang et al., 2017), alpha oscillations during resting state were specifically associated to the connectivity between mPFC and PCC (the anterior and posterior DMN). This specific connectivity plays a crucial role in self-relevant cognitive behaviors and choice justification (Tompson et al., 2016) and it is thought to mediate the sense of self (Washington and VanMeter, 2015). Interestingly, the antero-posterior DMN connectivity varies across human life span, it increases from childhood to adulthood (Washington and VanMeter, 2015) and it decreases during elderly (Vidal-Piñero et al., 2014; Andrews-Hanna et al., 2007), and an hyperconnectivity between mPFC/ACC and PCC has been correlated to neuropsychiatric disorders, e.g., schizophrenia and major depressive disorders (Whitfield-Gabrieli and Ford, 2012). Relative to the meditation practice, it has been proposed that a decrease in the anterior-posterior DMN connectivity could be linked to self-transcendence during meditation (Smigielski et al., 2019). However, how this specific connectivity is modulated after short meditation trainings in naïve practitioners has not been fully investigated yet.

Several studies revealed that short mindfulness meditation training programs can effectively increase health and well-being (Baer et al., 2012; Ditto et al., 2006; Harnett et al., 2010). A 1-week training of meditation promotes dispositional mindfulness, increases coping flexibility and reduces stress among novice meditators (Jones et al., 2019). Moreover, a few days of practice (4 days) can significantly reduce pain perception (Zeidan et al., 2011) and state anxiety (Zeidan et al., 2013) and even few minutes of mindful breathing reduced mind-wandering in a task of sustained attention as compared with passive relaxation or reading in naïve meditators (Mrazek et al., 2012). Nevertheless, the neural underpinnings accountable to the behavioral effects induced by short meditation trainings are still unknown. Given the neuroplastic nature of the antero-posterior DMN connectivity and its involvement in cognitive activity, e.g. self-referential processes (Tompson et al., 2016; Washington and VanMeter, 2015), specifically targeted by the meditation practice, we hypothesize that variations in this specific connectivity could in part underly behavioral changes induced by few days of meditation training in novices.

In this study, we aim to test the hypothesis that intracortical FC within the DMN, in particular between anterior and posterior DMN regions, is modulated after few days of meditation practice in novices. To address this question, we analyzed the effects of 6 days training of focused attention (FA) meditation on the breath in healthy novice meditators. We applied FA because this is one of the first practice generally introduced during the first weeks of mindfulness-based interventions (Kabat-Zinn et al., 1992; Segal et al., 2002). FA is a concentrative technique with a target object, which is usually the breath, that cultivates attentional control and monitoring skills (Cahn & Polich, 2006; Lutz et al., 2008) and it was reported to be specifically associated with deactivations in the PCC (Fox et al., 2016). Here, we recruited 32 healthy beginners, without any previous meditation experience, and we trained one group (n=17) for 6 consecutive days with FA on the breath (lasting 15 minutes), and a second group (n=15) with the same breath awareness practice but only on the first and last day of the experimental protocol. The DMN EEG functional connectivity was analyzed by using the exact low-resolution electromagnetic tomography software (eLORETA; Pascual-Marqui et al., 2002). For all participants, the brain activity was measured during both rest (trait) and FA (state) conditions before and after six days of training and was correlated with psychometric assessments of mindfulness.

## 2. Materials and methods

### 2.1. Participants

Thirty-two healthy volunteers (15 females) without any previous mindfulness experience or any other type of meditation practice took part in our study. All participants were Japanese undergraduate and graduate students, who have normal or corrected-to-normal visual acuity. Participants were randomly assigned to a group in which they performed the training of focused attention on the breath for six consecutive days (meditation group, MG, N=17) or to another group in which they performed two days of focused attention on the breath separated by 4 days of non-mindful activity (control group, CG, N=15). They were matched for sex, age and handedness between groups (Table 1). All participants gave written informed consent to participate in the study. After the completion of the study, participants were compensated for their time. The experiments were approved by the ethics committee of the School of Science and Technology, Meiji University, and conducted according to the principles and guidelines of the Declaration of Helsinki. Three subjects (2 from MG and 1 from CG) were excluded from the analysis because of EEG measurement errors.

**Table 1.** Subject’s demographics. Differences in sex and handedness were tested using Fisher’s exact tests. Differences in age were tested using the Mann-Whitney test.

|  | Control Group<br>N=14 | Meditation<br>Group N=15 | *P* |
| --- | --- | --- | --- |
| Gender (male/female) | 5/9 | 10/5 | 0.143 |
| Age (mean, standard deviation in parentheses) | 20.9(0.9) | 21.5(1.3) | 0.204 |
| Handedness (right/non-right) | 12/2 | 14/1 | >0.999 |

### 2.2. Focused attention on the breath protocol

At the beginning of the experiment on the first day, participants were introduced to the mindful practice by an experienced meditator and gave the instruction for the subsequent experiment. The mindful practice was based on focused attention (FA) on the breath, in which participants were given a commercially available guided audio with FA instructions recorded on a compact disc (Williams & Penman, 2016, “breath meditation”, CD Audio 1, Japanese translation). Each practice was composed as follow: one session of rest (rest condition) followed by 3 sessions of FA on the breath (meditation condition). Each session was 4 minutes and 20 seconds long and was separated by a few minutes of inter-session interval (Fig.1). During both rest and meditation conditions participants were enjoined to keep their eyes closed. During the rest condition subjects were instructed to rest, without engaging in any specific task or mental activity. The meditation group (MG) was asked to attend FA practices for 6 consecutive days, whereas the control group (CG) only on the first and the last day (day6). For all days of the experiment, FA practices were undertaken at the same hour of the day and same place (the authors’ laboratory) for each participant, in order to keep the same conditions as those for EEG recordings. EEG measurement and subjective questionnaire collection was conducted on the first and last day of the experiment for both groups. The experimenter followed up the MG throughout the entire week, to ensure that participants completed their practice. The CG was followed on the first and on the last day of the protocol and was asked to refrain from doing any meditation practice from day2 to day5. All subjects confirmed to have understood and successfully completed their practices.

**Figure 1.**
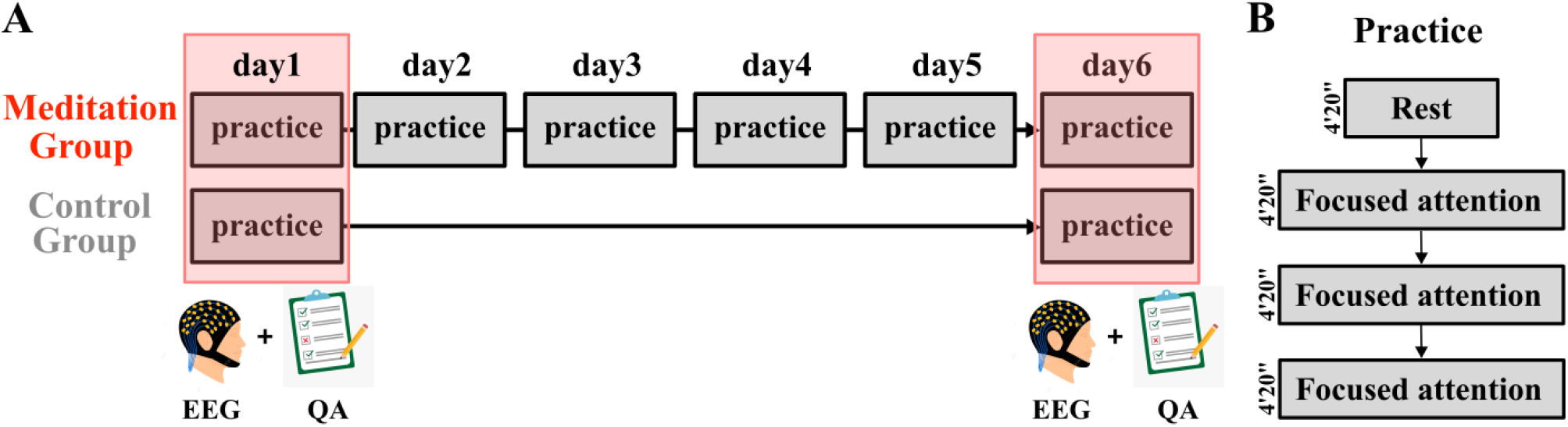
Experimental design. **A**. All participants underwent mindfulness assessments and EEG recordings at day1 and day6. The meditation group was required to engage in the FA practice for 6 consecutive days, whereas the control group only on day1 and day6. **B**. Each practice consisted of one session of rest followed by 3 consecutive sessions of FA. Each session lasted 4 minutes and 20 seconds and was separated by a few minutes of intersession interval. EEG recordings were undertaken both during rest and FA conditions. Abbreviations: FA, focused attention; QA, questionnaires; EEG, electroencephalography.

### 2.3. Self-reported measures

A battery of 4 standardized, mindfulness measures was given to all participants immediately after the FA practice on day1 and day6.

The *Five Facets Mindfulness Questionnaire* (FFMQ; Baer, 2006) is a 39-item constructed to assess five facets of mindfulness, including Observing, Describing, Acting with Awareness, Non-judging of Experience, and Non-reactivity to Experience. The measure uses a five-point Likert scale, ranging from 1 (*never or rarely true*) to 5 (*very often or always true*).

The short version *Freiburg Mindfulness Inventory* (FMI; Walach et al., 2006) is a 14-item measure recommended as more suitable for non-expert meditators. Each statement is evaluated using a four-point Likert scale ranging from 1 (*rarely*) to 4 (*almost always*) and assesses two facets of mindfulness: acceptance (e.g., “I accept unpleasant experiences“) and presence (e.g.,“I watch my feelings without getting lost in them“).

The *Mindful Attention Awareness Scale* (MAAS; Brown & Ryan, 2003; Fujino et al., 2015) is a self-report measure of mindfulness trait, consisting of 15-items assessing the inclination to be mindful of present experiences. The measure uses a 6-point Likert scale, ranging from 1 (*almost always*) to 6 (*almost never*). Higher total scores on the MAAS indicate greater levels of dispositional mindfulness.

The *Toronto Mindfulness Scale* (TMS; Lau et al., 2006) is a 13-item self-report scale designed to measure the subjective experience of a mindfulness state in reference to an immediately preceding mindfulness meditation practice. It includes total score and two sub-scales: *Curiosity* and *Decentering*. *Curiosity* reflects “awareness of present moment experience with a quality of curiosity”; *Decentering* represents “awareness of one’s experience with some distance and disidentification rather than being carried away by one’s thoughts and feelings” (Lau et al., 2006). Items described what subjects have just experienced on a 5-point Likert scale from 0 (*not at all*) to 4 (*very much*). While FFMQ, FMI and MAAS questionnaires measure dispositional mindfulness (trait mindfulness), the TMS scale measures state mindfulness.

### 2.4. EEG recordings

EEG signals (g.USBamp, g.tec Inc., Schiedlberg, Austria) were recorded from 30 scalp sites (Fp1, Fp2, F7, F3, Fz, F4, F8, FT7, FC3, FCz, FC4, FT8, T7, C3, Cz, C4, T8, TP7, CP3, CPz, CP4, TP8, P7, P3, Pz, P4, P8, POz, O1 and O2), located according to the extended international 10/20 system. An electrode placed on the left ear lobe was used as a reference. The vertical electrooculography (EOG) was recorded from electrodes placed above and below the left eye to monitor blinks or vertical eye movements. EEG and EOG were amplified and digitized at 512 Hz with a band-pass filter of 0.5 to 100 Hz. The subjects were instructed to refrain from opening their eyes and moving their bodies during EEG recording.

### 2.5. EEG data Analysis

Brain activity of all subjects was recorded through electroencephalography (EEG) during both rest and meditation conditions on day1 and day6. All the EEG data analyses were performed with MATLAB R2016b (MathWorks Inc., Natick, MA, USA) environment. First, we preprocessed EEG signals using MATLAB toolbox EEGLAB version 14.1.2b (Delorme & Makeig, 2004). We performed down sampling process for the EEG signals from 512 to 128 Hz and applied a band-pass filter from 1 to 60 Hz to the EEG signals. Alternating current line noise was removed using EEGLAB. An epoch of 264 sec, starting 5 sec before the onset, was selected from EEG signal (i.e., −5 to 259 sec from the onset). Then, we conducted infomax independent components analysis (ICA) to reduce or eliminate artifacts from blinks or eye movement.

We used standardized low-resolution brain electromagnetic tomography (LORETA) (Pascual-Marqui, 2002) software to compute the cortical three-dimensional distribution of electrical neuronal activity. In particular, we applied exact LORETA (eLORETA), which provides exact localization with zero error and produces results with more accuracy compared to standardized LORETA (sLORETA) (Jatoi et al., 2019). The detailed description of the method can be found in (Pascual-Marqui, 2002, 2007a). For a correct positioning of electrodes at the surface of the head (important requisite for the stability of the source localization) we first measured the locations of electrodes for each subject using a 3D magnetic space digitizer (FASTRAK, Polhemus, VT, USA) and, second, we implemented the real coordinates in eLORETA which enables to fit the positions of the electrodes onto the coordinates of the Montreal Neurological Institute average MNI brain map (MNI 152; Mazziotta et al., 2001) with a 12-parameter affine transformation that finally produces coordinates that really lie on the MNI152 scalp. Third, by using the transformation matrix computed from these coordinates, EEGs data for each subject were transformed into eLORETA files.

In order to investigate the effects of six days of FA on the breath on the DMN-FC, we computed functional connectivity analysis on a priori selected regions of interests (ROIs) as components of the DMN. To create the ROIs, we used eLORETA software which defined the MNI coordinates of the cortical voxels underlying the electrode sites (Pascual-Marqui et al., 2011). In particular, the signal at each cortical ROI consisted of the average electric neuronal activities of all voxels belonging to that ROI, as computed with eLORETA. As illustrated in Fig.2 and Table2, the ROIs were composed as follows: left (ROI1) and right (ROI2) superior frontal gyrus (SFG); left (ROI3) and right (ROI4) ventral/dorsal anterior cingulate cortex (v/dACC) and ventromedial prefrontal cortex (vmPFC); left (ROI5) and right (ROI6) posterior cingulate cortex (PCC) and precuneus (PCUN). These ROIs were chosen based on their involvement in mindfulness meditation and the DMN (Brewer et al., 2011; Doll et al., 2015) and on previous studies investigating the DMN with EEG (Hata et al., 2016; Imperatori et al., 2016; Thatcher et al., 2014). The connectivity analysis was performed by the computation of a measure of linear dependence (coherence), which provides a measure of linear similarity between signals in the frequency domain (Pascual-Marqui, 2007b; Pascual-Marqui et al., 2011). The algorithm is a measure of true physiological connectivity not affected by volume conduction and low spatial resolution, as shown in Pascual-Marqui et al. (2011). Intracortical coherence has been widely used elsewhere (Lehmann et al., 2014; Painold et al., 2020) and even applied in meditation studies (Lehmann et al., 2012; Milz et al., 2014). Here, we measure intracortical coherence between all possible 16 pairs of the 6 ROIs for 3 independent EEG frequency bands based on their association with meditation and DMN (Berkovich-Ohana et al., 2014; Faber et al., 2004; Fingelkurts et al., 2016; Lee et al., 2018a): theta (4-8Hz), alpha (8-13Hz) and gamma (30-60Hz).

**Figure 2.**
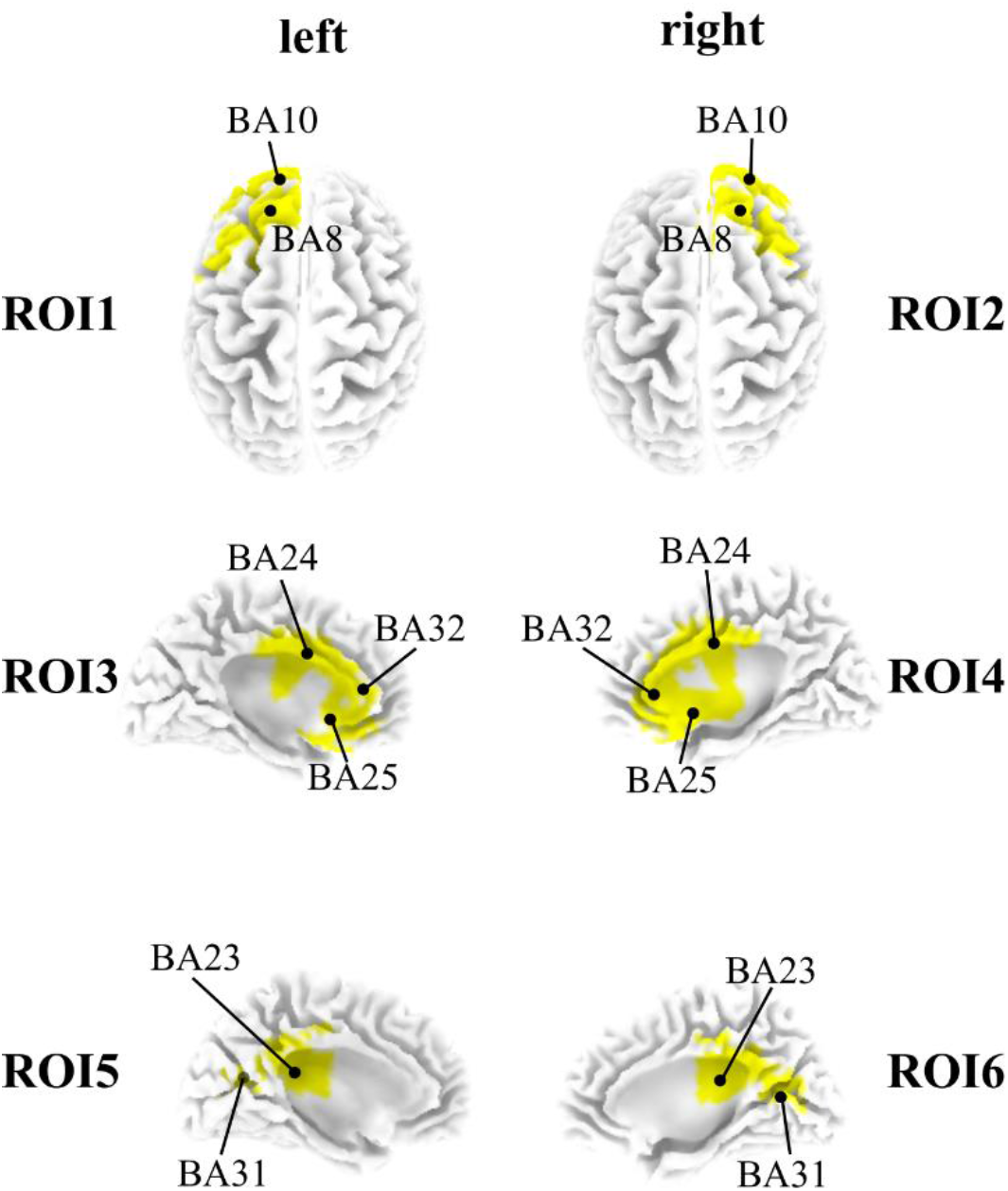
DMN selected regions of interests (ROI). Anatomical locations of the 6 ROIs used in the connectivity analysis. The description for the anatomical regions and Brodmann areas included in each ROI are described in Table 2.

**Table 2.** List of anatomical regions of interest (MNI coordinates at ROI centroid).

| ROI | MNI coordinates |  |  | Lobe | Anatomical region | Abbreviation | BA |
| --- | --- | --- | --- | --- | --- | --- | --- |
| 1 | -20 | 30 | 50 | Frontal Lobe | superior frontal gyrus | SFG | 8 |
| 1 | -25 | 55 | 5 | Frontal Lobe | superior frontal gyrus | SFG | 10 |
| 2 | 20 | 25 | 50 | Frontal Lobe | superior frontal gyrus | SFG | 8 |
| 2 | 25 | 55 | 5 | Frontal Lobe | superior frontal gyrus | SFG | 10 |
| 3 | -5 | 0 | 35 | Limbic Lobe | ventral anterior cingulate | vACC | 24 |
| 3 | -10 | 20 | -15 | Frontal Lobe | ventromedial prefrontal cortex | vmPFC | 25 |
| 3 | -5 | 30 | 20 | Limbic Lobe | dorsal anterior cingulate | dACC | 32 |
| 4 | 5 | 0 | 35 | Limbic Lobe | ventral anterior cingulate | vACC | 24 |
| 4 | 5 | 15 | -15 | Frontal Lobe | ventromedial prefrontal cortex | vmPFC | 25 |
| 4 | 5 | 30 | 20 | Limbic Lobe | dorsal anterior cingulate | dACC | 32 |
| 5 | -5 | -40 | 25 | Limbic Lobe | posterior cingulate | PCC | 23 |
| 5 | -10 | -50 | 30 | Parietal Lobe | precuneus | PCUN | 31 |
| 6 | 5 | -45 | 25 | Limbic Lobe | posterior cingulate | PCC | 23 |
| 6 | 10 | -50 | 35 | Parietal Lobe | precuneus | PCUN | 31 |

### 2.6. Statistics

#### 2.6.1. Questionnaires

The data were checked for normality with the Shapiro-Wilk Test. A repeated measure ANOVA was then performed for each questionnaire scale and subscale, with group (meditation and control groups) as between-subjects factor and time-point (day1 and day6) as within-subjects factor. Statistical significance was evaluated at p<0.05. Bonferroni correction was applied for multiple comparisons.

#### 2.6.2. EEG Functional Connectivity

To test whether functional connectivity changed following six days of FA, a two-way ANOVA was applied on the z-transformed day6-day1 variations (day6 minus day1), for each frequency bands and each connectivity combination with group (meditation and control groups) and condition (rest and FA) as factors. Bonferroni correction was applied for multiple comparisons.

#### 2.6.3. Correlation

To check whether the change in EEG functional connectivity cooccurred with the change in subjective questionnaire score, we further calculate the correlation between those. A change in questionnaire score was calculated by subtracting the data on day1 from that on day6 (*day6 – day1*). For functional connectivity changes, the data for 3 FA sessions were averaged and normalized with the data of rest session *(FA/rest)*, and the change was calculated by subtracting day1 from day6 scores: *(FA/rest)day6 – (FA/rest)day1*. The Spearman’s rank correlation (r_s_) between questionnaire scores and EEG functional connectivity data was used at each frequency bands and for all connectivity combinations, as these data did not fit the normality assumption. Bonferroni correction was applied for multiple comparisons.

## 3. Results

### 3.1. Questionnaires data

Repeated measures ANOVAs confirmed significant effect of day for one of the four mindfulness scales, namely TMS (TMS: F_(1,27)_ = 7.493, P = 0.0108; FFMQ: F_(1,27)_ = 0.0604. P = 0.8077; FMI: F_(1,27)_ = 3.213, P =0.0843; MAAS: F_(1,27)_ = 0.4002, P = 0.5323; Table 3). Subsequent Bonferroni’s multiple comparisons tests confirmed significant increases in the MG (TMS: t_(15)_ = 2.736, P = 0.0217), but not in the CG (TMS: t_(14)_ = 1.163, P = 0.5099; Table 3). In particular, repeated measures ANOVA on the two TMS subscales confirmed significant effect of day for both curiosity and decentering subscales (curiosity: F_(1,27)_ = 5.332, P= 0.0288; decentering: F_(1,27)_ = 9.524; P = 0.0046). Subsequent Bonferroni’s multiple comparisons tests confirmed significant increase in the MG (t_(15)_ = 3.227, P = 0.0065), but not in the CG (t_(14)_ = 1.174, P = 0.5017; Table3) selectively in the decentering subscale, while no differences were found in the curiosity subscale (MG: t_(15)_ = 1.562, P = 0.2599; CG: t_(14)_ = 1.702, P = 0.2006; Table 3). That is, six consecutive days of FA on the breath significantly increase the mindfulness scores on the TMS scale, in particular on the decentering subscale.

**Table 3.** Averaged questionnaires scores (± standard error) and relative *t*- and *p*-values before (day1) and after (day6) six days of FA on the breath. Bonferroni’s multiple comparisons test: day1 vs day6, *p<0.05, **p<0.01.

|  | Meditation group |  |  |  | Control group |  |  |  |
| --- | --- | --- | --- | --- | --- | --- | --- | --- |
| | day1 | day6 | $t_{15}$ | $p$ | day1 | day6 | $t_{14}$ | $p$ |
| **TMS** | **25.47(2.0)** | **29.2(2.2)\*** | **2.736** | **0.0217** | 26.36(1.9) | 28(0.4) | 1.163 | 0.5099 |
| curiosity | 11.47(1.3) | 12.73(1.2) | 1.562 | 0.2599 | 11.64(1.4) | 13.07(1.0) | 1.702 | 0.2006 |
| decentering | **14.00(0.9)** | **16.47(1.1)\*\*** | **3.227** | **0.0065** | 14.71(0.9) | 15.64(1.0) | 1.174 | 0.5017 |
| **FMI** | 35.47(1.3) | 37.73(1.7) | 1.680 | 0.209 | 36.57(1.5) | 37.78(1.3) | 0.869 | 0.7845 |
| **MAAS** | 3.59(0.1) | 3.83(0.1) | 1.879 | 0.1422 | 3.70(0.1) | 3.58(0.2) | 0.935 | 0.7157 |
| **FFMQ** | 118.8(4.4) | 119.6(3.2) | 0.194 | >0.999 | 121.21(4.2) | 119.15(4.8) | 0.529 | >0.999 |
| observing | 22.73(1.7) | 21.86(1.3) | 0.961 | 0.6899 | 21.28(1.4) | 21.78(1.1) | 1.607 | 0.2393 |
| non reaction | 22.2(1.8) | 21.47(1.1) | 0.752 | 0.9169 | 21.28(0.9) | 20.28(0.9) | 0.991 | 0.6610 |
| non judgement | 24.53(1.7) | 23.87(1.3) | 0.473 | >0.999 | 23.71(0.9) | 25.35(1.4) | 1.126 | 0.5401 |
| describe | 21.73(1.5) | 22.47(1.5) | 0.502 | >0.999 | 22.21(1.5) | 22.36(1.4) | 0.094 | >0.999 |
| action with awareness | 27.6(2.6) | 29.87(2.2) | 1.709 | 0.1977 | 30.71(3.1) | 29.36(3.1) | 0.989 | 0.6631 |

### 3.2. Functional connectivity changes in the DMN

We compared functional connectivity changes from day1 to day6 (day6 minus day1) in the two groups for each connectivity combination and for each frequency band. Two-way ANOVAs confirmed a significant main effect of group in three of the twelve ROI connectivity combinations in theta (ROI3-ROI5: t_(1,112)_ = 4.42, P = 0.0377; ROI3-ROI6: t_(1,112)_ = 8.505, P = 0.0043; ROI4-ROI6: t_(1,112)_ = 5.28, P = 0.0234), in four connectivity in alpha (ROI3-ROI5: t_(1,112)_ = 5.92, P = 0.0166; ROI3-ROI6: t_(1,112)_ = 9.047, P = 0.0032; ROI4-ROI5: t_(1,112)_ = 4.36, P = 0.0391; ROI4-ROI6: t_(1,112)_ = 4.63, P = 0.0335), and in one connectivity in gamma (ROI3-ROI6: t_(1,112)_ = 5.02, P = 0.0270). Subsequent Bonferroni’s multiple comparisons tests confirmed significant decrease in the MG compared to the CG selectively in the meditation condition for the connectivity between ROI3 and ROI6 in two frequency bands (theta: t_(112)_ = 3.051, P = 0.0029; alpha: t_(112)_ = 3.054, P = 0.0028; Fig.3A, B, D), whereas no differences were found in gamma band (t_(112)_ = 2.45, P = 0.095; Fig.3C), neither in the other connectivity analyzed (p>0.05). No differences were found between the two groups in the rest condition (theta: t_(112)_ = 1.606, P = 0.111; alpha: t_(112)_ = 1.71, P = 0.09; gamma: t_(112)_ = 1.173, P > 0.999). Overall, the change in functional connectivity in the MG is significantly lower than that in the CG, in theta and alpha bands.

**Figure 3.**
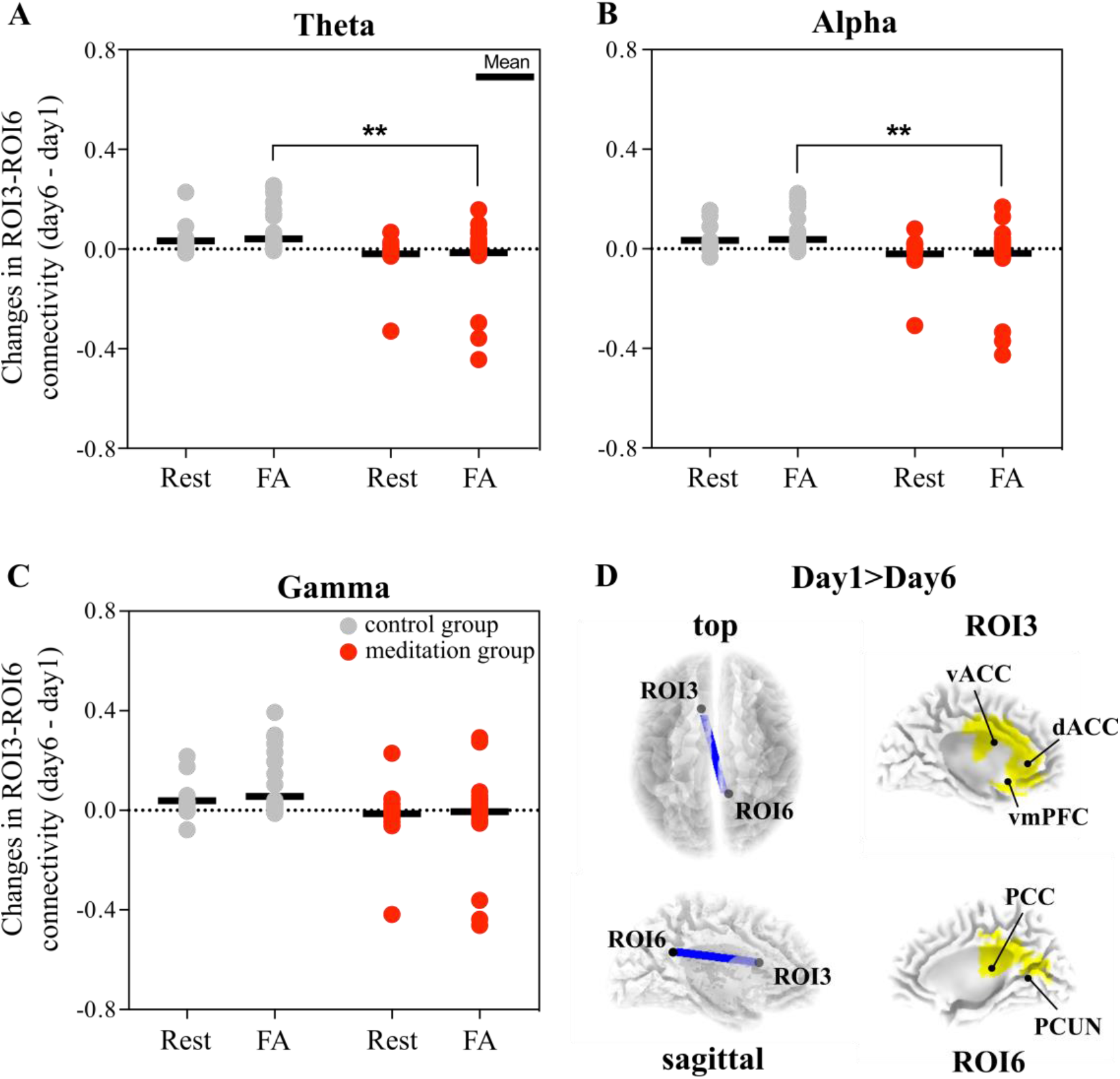
Connectivity changes before and after 6 days of FA on the breath. Panels A-C show day6 vs day1 variation in connectivity strength between ROI3 and ROI6 in meditation (red) and control (grey) groups, in theta (A), alpha (B) and gamma (C) bands. Panel D shows the connectivity between the ROI3 (left vACC, dACC and vmPFC) and ROI6 (PCC and PCUN). Bonferroni’s multiple comparisons test: day1 vs day6, *p<0.05, **p<0.01. Abbreviations: FA, focused attention (on the breath); vACC, ventral anterior cingulate cortex; dACC, dorsal anterior cingulate cortex; vmPFC, ventromedial prefrontal cortex; PCC, posterior cingulate cortex; PCUN, precuneus.

### 3.3. Negative correlation between the TMS decentering scale and functional connectivity changes

A correlation analysis was used to investigate whether the changes in questionnaire scores (day6 minus day1) were associated with functional connectivity changes during the meditation condition. In the MG, we found a negative correlation between changes in TMS decentering scores and relative changes in ROI3-ROI6 connectivity in both theta (r_s_ = −0.78, P = 0.0006) and alpha bands (r_s_ = −0.69, P = 0.0041; Fig.4A, B and Table 4), while the same correlation in gamma band didn’t reach statistical significance after correcting for multiple comparisons (r_s_ = −0.56, P = 0.0292; Table 4). No significant correlations were found in the CG (theta: r_s_ = −0.01, P = 0.9701; alpha: rs = 0.19, P = 0.5111; gamma: rs = 0.15, P = 0.0361; Fig. 4A, B and Table 4). The correlation between TMS decentering scale with other connectivity analyzed didn’t show statistical significance (P > 0.05 after correcting for multiple comparisons), neither by considering TMS total scores nor TMS curiosity subscale scores (P > 0.05 after correcting for multiple comparisons). For the sake of completeness, we also checked correlations between all mindfulness questionnaires scores analyzed in section 3.1 with connectivity variations, but we didn’t find any statistical significance (P>0.05). Overall, these data show that after 6 consecutive days of FA on the breath, the higher the TMS decentering score is, the lower the functional connectivity between left d-vACC / vmPFC and the right PCC/PCUN is.

**Figure 4.**
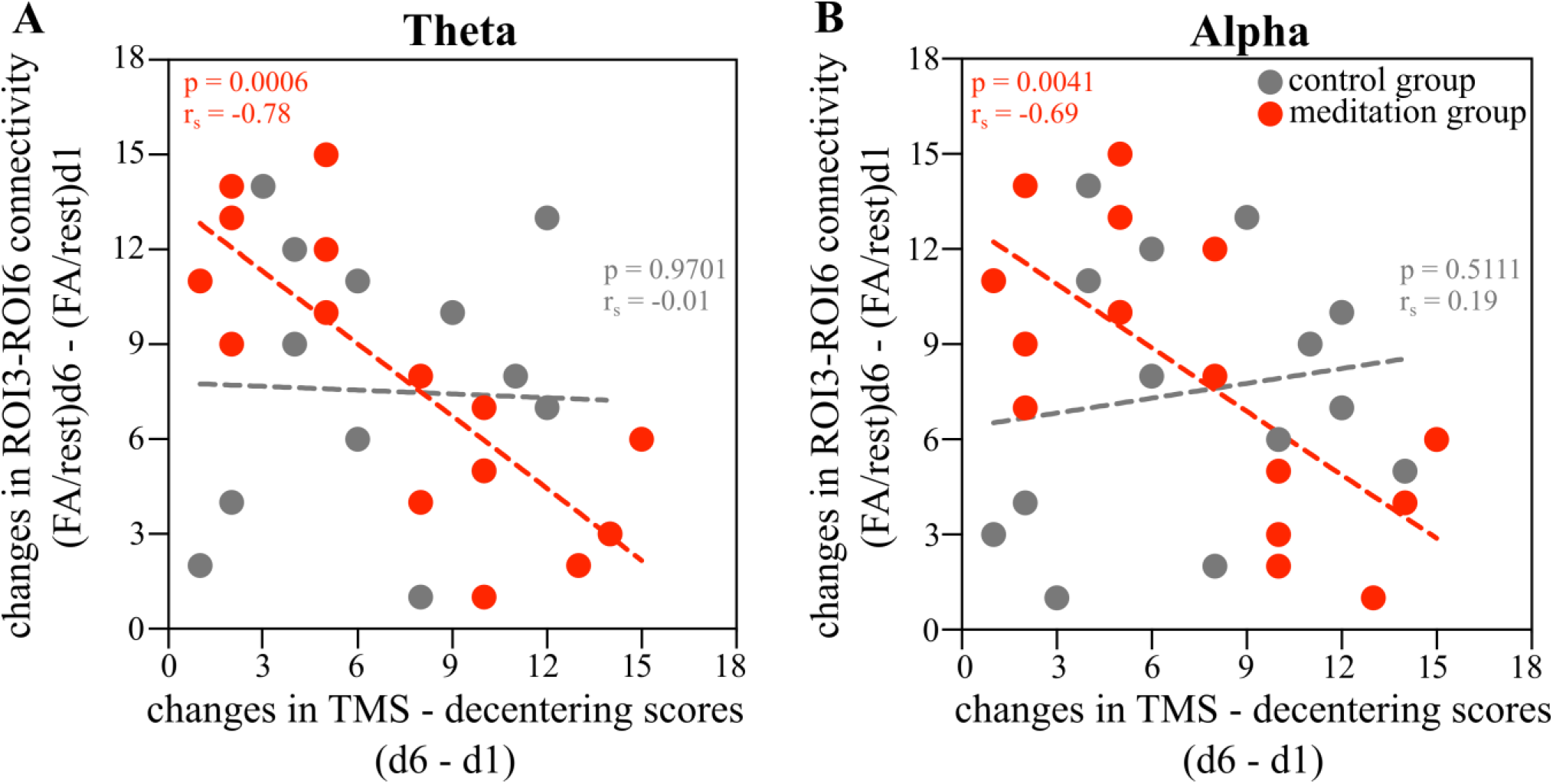
Correlation between day6 – day1 variations in TMS decentering scores and ROI3-ROI6 functional connectivity. **A, B.** Scatterplots describe the correlation between day6-day1 variations in TMS decentering scores and connectivity strength during meditation condition between ROI3 and ROI6 in theta (A) and alpha (B) bands in meditation (red) and control (grey) groups. Bonferroni corrected for 12 connectivity combinations with corrected *p*-value, *p*<0.0042.

**Table 4.**
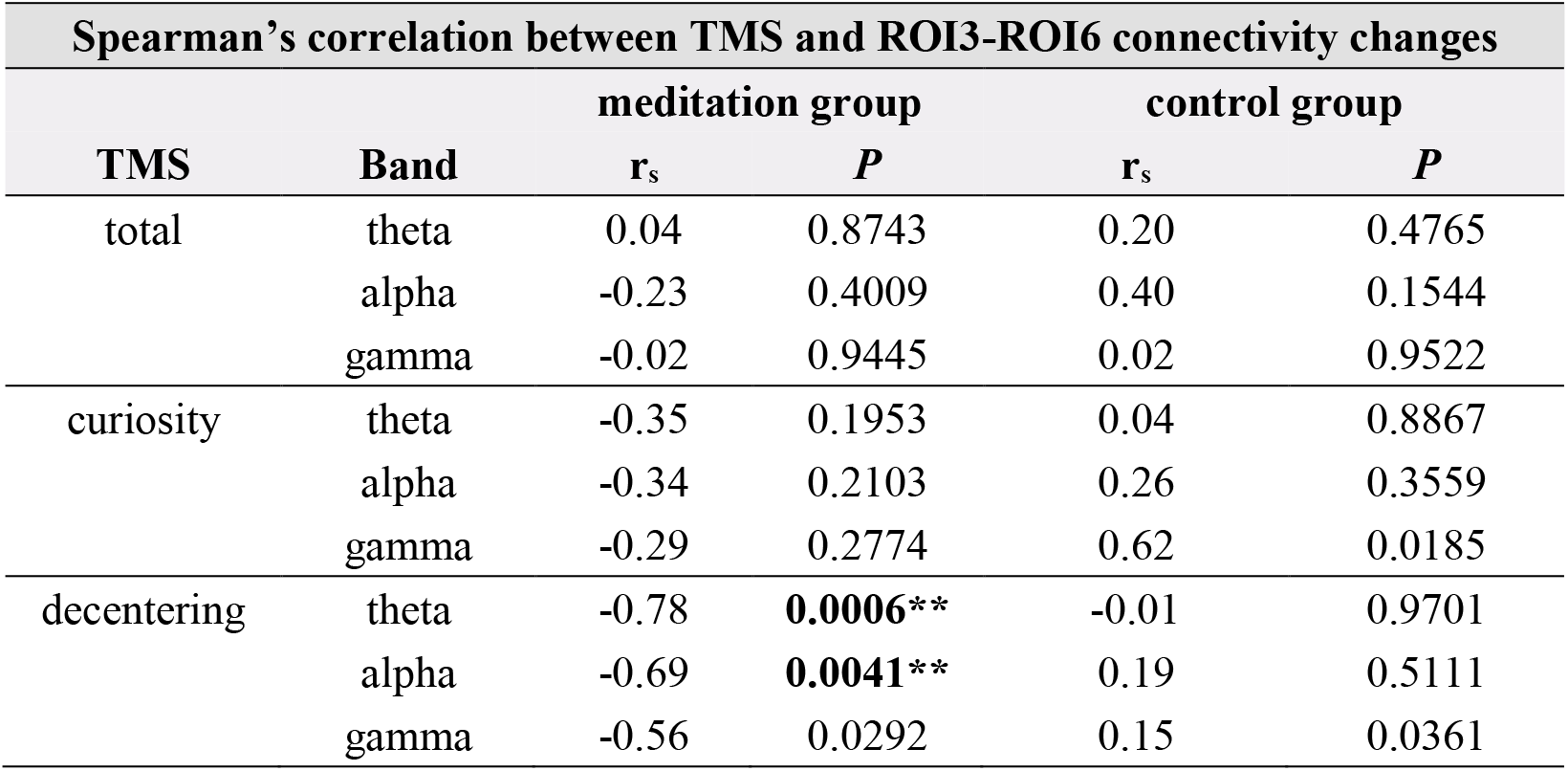
Correlation between TMS scores and ROI3-ROI6 variations from day1 to day6. Correlation between TMS variations of total scores, curiosity and decentering subscales with functional connectivity variation during meditation condition for MG (left) and CG (right) in theta, alpha and gamma bands. Significant p-values were considered only when p<0.0042**. Bonferroni correction for 12 connectivity combinations. Abbreviations: rs, Spearman correlation coefficient; P, p-value.

| Spearman's correlation between TMS and ROI3-ROI6 connectivity changes |  |  |  |  |  |
| --- | --- | --- | --- | --- | --- |
| TMS | Band | meditation group |  | control group |  |
| | | $r_s$ | $P$ | $r_s$ | $P$ |
| total | theta | 0.04 | 0.8743 | 0.20 | 0.4765 |
|  | alpha | -0.23 | 0.4009 | 0.40 | 0.1544 |
|  | gamma | -0.02 | 0.9445 | 0.02 | 0.9522 |
| curiosity | theta | -0.35 | 0.1953 | 0.04 | 0.8867 |
|  | alpha | -0.34 | 0.2103 | 0.26 | 0.3559 |
|  | gamma | -0.29 | 0.2774 | 0.62 | 0.0185 |
| decentering | theta | -0.78 | **0.0006\*\*** | -0.01 | 0.9701 |
|  | alpha | -0.69 | **0.0041\*\*** | 0.19 | 0.5111 |
|  | gamma | -0.56 | 0.0292 | 0.15 | 0.0361 |

## 4. Discussion

The aim of this study was to investigate the effects of few days of focus attention meditation on the breath on the functional connectivity between anterior and posterior DMN regions. We found that 6 days meditation training significantly increased the mindfulness state (higher scores in the TMS scale), in particular, the decentering state. We also showed that the difference in FC before and after the training in the meditation group (MG) between anterior-posterior DMN regions was significantly lower compared to the control group (CG). The functional connectivity change between the left ACC/vmPFC and the right PCC/PCUN was evinced for EEG theta and alpha bands during the meditation condition, whereas no differences were found during the rest condition. Finally, participants’ scores on the TMS decentering scale negatively correlated with the amount of functional connectivity difference before and after the training. Overall, our results suggest that even a few days of meditation training in novice practitioners can lead to an increase in psychometric assessments of mindfulness state and to a modulation of the anterior-posterior DMN functional connectivity.

### 4.1. Brief meditation training improves a decentering state

In order to investigate which aspects of the mindful experience are influenced by a short FA practice, we used 4 different mindfulness questionnaires (FFMQ, FMI, MAAS and TMS), which cover several aspects of the mindfulness experience. Based on previous reports, an increase in self-reported mindfulness scores will be the outcome of an increase in mindfulness (Bohlmeijer et al., 2011; Chambers et al., 2008; Lau et al., 2006). Here, we obtained a significant improvement in the TMS scale, in particular, on the *decentering* subscale (Table 3). This result is coherent with the state-like nature of the TMS scale (Baer, 2011; Lester & Murrell, 2019), as it is the case for the State Mindfulness Scale (Lester & Murrell, 2019; Tanay & Bernstein, 2013), meaning that these questionnaires refer to the practice of mindfulness *per se*, evaluating a transitory effect related to the previous mindfulness practice. Theses scales have been validated with naïve meditators after short meditation training (Anderson et al., 2007; Gayner et al., 2012; Lau et al., 2006; Tanay & Bernstein, 2013). On the other hand, the other questionnaires used in our study conceived mindfulness as a dispositional, trait-like variable which is consistent over time and across situations (continuing in everyday life, beyond the practice) (Baer et al., 2006), so that they are more appropriate for experienced meditators and/or longer mindfulness interventions.

TMS results further revealed that the psychological traits evoked by six days of focused attention on the breath specifically promote a decentering state, a shift from identifying personally with thoughts and feelings in relation to one’s experience in a wider field of awareness (Lau et al., 2006). The decentering factor is akin the concept of “reperceiving”, and it involves a “shift in perspective”, important indicator of mindfulness. Goleman (1980) stated that “The first realization in ‘meditation’ is that the phenomena contemplated are distinct from the mind contemplating them”. Supporting this claim, we observe that “decentering” is one of the first factors appearing at the initial stages of mindfulness. By contrast, a six-day FA training may not be sufficient in increasing curiosity of one’s experiences, probably because a curiosity attitude may emerge after a decentering state is reached. In other words, decentering from or disidentification with experience may be a necessary prerequisite for cultivating curiosity.

### 4.2. A specific anterior-posterior DMN connectivity is reduced after six days of FA in theta and alpha bands

DMN is often reported with its modulations on activity and connectivity induced by the meditation practice, displayed in particular in experienced meditators (Berkovich-Ohana et al., 2014; Brewer et al., 2011; Garrison et al., 2015; Pagnoni et al., 2008). To the best of our knowledge, our study is the first report that explores the DMN connectivity in novices at the beginning of the mindfulness training involving the focus attention on the breath. In particular, we focused on the anterior-posterior DMN functional connectivity because it has been shown to play a crucial role in self-relevant cognitive behaviors (Tompson et al., 2016). The result showed that the functional connectivity between the left dvACC/vmPFC with the right PCC/PCUN showed a significant reduction in the MG compared to the CG. Those regions are considered central nodes of the DMN (Buckner et al., 2008). The ACC is associated with attention regulation (Pardo et al., 1990) and emotion (Bush et al., 2000) and neuroplastic changes related to the meditation practice has been documented in this region (Grant et al., 2010; Hölzel et al., 2007; Manna et al., 2010). The vmPFC is involved in self-referential processing (Andrews-Hanna et al., 2010; Gusnard et al., 2001) and in the top-down regulation of emotional responses (Davidson, 2002; Phelps et al., 2004) thanks to its dense projection to the amygdala (Amaral et al., 1992). The precuneus has been proposed to be involved in the neural correlates of self-consciousness, in particular in the self-related mental representations during rest (Cavanna & Trimble, 2006). Finally, the PCC is considered as a cortical hub, since it has dense structural connectivity to widespread brain regions (Leech et al., 2012). The PCC appears to be involved in internally directed thoughts (e.g., memory recollection), plays an active role in the control of cognition, and interacts with frontoparietal networks to regulate the balance between internally and externally directed cognition (Leech et al., 2012). Brewer and Garrison (2014) proposed that the PCC is related to disentanglement from self-referential processing. Considering the relation of these brain regions with internal mental attention, and in view of our findings, we can speculate that few days of mindful attention toward an internal state (in this case the breath) can reduce the amount of self-referential mind wandering. Self-directed thoughts with an unfocused state of mind is typical of mind-wandering, thoughts that can predict negative emotions and pessimistic mental states (Killingsworth & Gilbert, 2010), which lead to negative loops difficult to break. A higher DMN activity and connectivity has been associated with mind wandering (Christoff et al., 2009; Mason et al., 2007).

Several studies reported that the reduction of mind wandering is considered one of the most beneficial effects of focused attention meditation in non-experts (Morrison et al., 2014; Mrazek et al., 2012; Rahl et al., 2017) which may lead in turn to cognitive regulation and metacognition (Kerr et al., 2013). In beginners, FA meditation mainly involves disengagement from mind-wandering, and thus deactivation and reduced connectivity of the DMN. Accordingly, it has been shown that even few minutes of mindfulness practice can reduce mind wandering (Mrazek et al., 2012). Our results suggest that even 6 days of FA training, through a reduced connectivity within anterior-posterior DMN regions, can reduce mind-wandering. In a previous study (Scheibner et al., 2017), it has been shown that after 5 days of internal or external attention meditation (for a total amount of 3 hours) in beginners, regions associated with the DMN (e.g., mPFC and PCC) were less activated during the mindful attention condition compared to a mind-wandering phase. This result is interesting in that brain activity was detected during a mindful state, where participants were asked to report if their mind was wandering during the focus attention meditation, and not during a rest phase. Accordingly, we also found a reduced connectivity within the DMN during the focused attention meditation condition, while no differences were found during the rest condition. This result advises that 6 days of FA meditation on the breath induce changes in mindfulness *state* (temporarily changes), rather than changes in mindfulness *trait* (changes in personality that may arise after a longer period of practice).

Intriguingly, here we found that EEG theta and alpha bands showed modifications after the meditation training, whereas no statistically significant alterations were found in gamma band. Understanding these findings in relation to previous electrophysiological correlates of DMN in relation to meditation is difficult because the literature is rather inconsistent. Previous studies reported either increases (Berkovich-Ohana et al., 2014; Faber et al., 2004; Lee et al., 2018b) or decreases (Berkovich-Ohana et al., 2014; Faber et al., 2004; Fingelkurts et al., 2016; Lehmann et al., 2012) depending on the frequency band analyzed (from delta to gamma), on the condition compared (rest, meditation or active task), on the meditation practice involved (focused attention, open monitoring or transcendental meditation) and on the expertise of the meditator (novices or relative experts). Activity on the high frequency range (gamma) has been observed in the earlier studies on meditation (Banquet, 1973; Das et al., 1957; Lutz et al., 2008). The overall reduction, rather than specific to the DMN, in gamma EEG-FC was reported during the transition from resting state to an active task (Berkovich-Ohana et al., 2014) or during meditation compared to rest (Faber et al., 2004) in experienced meditators. On the other hand, studies focusing on naïve practitioners before and after canonical mindfulness programs (e.g., MBSR) or shorter meditation protocols showed modulations of the theta and alpha bands following the training, while they did not report alterations of gamma bands (Lee et al., 2018b; Rodriguez-larios et al., 2020) In accordance with the later studies, we didn’t find modulation in gamma bands but only in theta and alpha bands.

Theta activity has been linked to various type of cognitive activity, including switching and orienting attention (Dietl et al., 1999), but data evaluating changes in this specific frequency band during a focus attention meditation task, in particular in relation to the DMN regions, are still lacking. By contrast, more clues related the alpha band. These oscillations dominate EEG of humans in the absence of external stimuli when internal life (mind wandering and spontaneous thoughts) is most pronounced (Palva & Palva, 2007; Shaw, 2003) and increased alpha power has been linked to an internally oriented state of attention. Therefore, modulation in alpha frequency band would reflect a modification in internally oriented attention. Considering that both alpha (Fingelkurts et al., 2016; Lee et al., 2018a; Lee et al., 2018b; Rodriguez-larios et al., 2020) and theta (Jacobs & Friedman, 2004; Lee et al., 2018a; Lee et al., 2018b; Rodriguez-larios et al., 2020) bands are altered by meditation practice, regardless of the expertise of the meditators, we hypothesize that alterations on these frequency bands may be regarded as a first basic change in the course of meditative development in the brain. In other words, these first primary alterations should be easily accessible even by inexperienced practitioners within a short training period.

### 4.3. Relationship between mindfulness assessments and DMN connectivity

In the MG, we found that the change in the decentering state was increased (higher TMS decentering scores) after the 6-day meditation training as the change in the functional connectivity within DMN regions decreased. This result represents individual differences that may emerge during the early stage of meditation training. In other words, the more the participant undergoes decentering from their proper thoughts, the lower the anterior-posterior DMN functional connectivity becomes. A similar result was found by Xiao and colleagues (2019), in which they showed that 8 weeks of MBSR training leads to a significant negative correlation between the FFMQ scale (in particular the observing component) and regional homogeneity values (an indicator of functional connectivity) within the left anterior ACC. Even though they investigated a longer meditation protocol (eight weeks) compared to our protocol (six days), it is interesting in that they also found a reduced functional connectivity in crucial regions within the DMN as a function of an improvement of a self-reported mindfulness scale. This result is also consistent with the study of Doll and coworkers (2015), in which they found that after 2 weeks of mindful attention on the breath, the connectivity of the anterior and the posterior parts of the DMN is negatively correlated to MAAS scores. We did not find any correlation with neither the FFMQ nor the MAAS scores, probably due to the *trait-like* nature of these 2 scales in contrast to the *state-like* nature of the TMS scale, as already discussed in section 4.1. Indeed, both the aforementioned studies (Doll et al., 2015; Xiao et al., 2019) reported negative correlation with FFMQ or MAAS with resting-state functional MRI, which is considered to ascribe to changes in mindfulness trait. By contrast, our finding of EEG functional connectivity changes in anterior-posterior DMN regions is considered to reflect the variation in mindfulness state during meditation.

### 4.4. Limitations

While our results offer important highlights on the neural and subjective effects induced by a short training of focused attention on the breath, there are some limitations that must be considered. Firstly, the self-reported measures of mindfulness implied here are not exhaustive in detecting an objective *state* or *trait* of mindfulness in participants, since they are subjective measures. Even though EEG measures can be intended as objective measures for mindfulness, it will be important also to encourage additional behavioral observations for the evaluation of mindfulness experience. Secondly, the time of our intervention was shorter (six days) than common mindfulness interventions (e.g., MBSR). Therefore, it is possible that, by prolonging the meditation training after six days, we will be able to find additional significant differences, but this would go beyond the purpose of the present study. Indeed, reporting that even the first week of a FA training leads to positive neuronal effects could prompt people to engage in such practices more easily. Accordingly, a previous study (Lee et al., 2018b) has demonstrated that only four weeks of an online-mind-body training can reduce anxiety and anger traits in women. Thus, it will be interesting to investigate the psychological effects (e.g., reducing anxiety and/or stress) of short and specific meditation interventions (e.g., FA on the breath and/or sounds, open monitoring or loving kindness). Thirdly, the analysis with LORETA may have methodological limitations when exploring brain function, because of its low spatial resolution. EEG-source imaging techniques provide very good temporal resolution and they have extensively applied for the analysis of DMN functional connectivity (Thatcher et al., 2014; Imperatori et al., 2016, 2019; Farina et al., 2017), but their accuracy strongly depend by the density (higher with high-density EEGs) and disposition of the electrodes array. However, LORETA source modelling with low-density EEGs has been validated with good results in simultaneous fMRI/PET-EEG (Horacek et al., 2007; Cannon et al., 2012; Zotev et al., 2020) and also applied with 30 channel EEGs (Thatcher et al., 2014; Travis et al., 2017; Farina et al., 2017; Lee et al., 2018; Imperatori et al., 2016, 2019) as well as with 19 channels in clinical studies (Hata et al., 2016; Lanzone et al., 2020). Here, in order to overcome some critical issues of this method, we used some pre-requisite parameters for conducting functional connectivity on EEG source estimates, such as measuring real electrodes coordinates with a 3D digitizer and applying exact LORETA (eLORETA) with less localization error and clear visibility compared to standardized LORETA (JJatoi et al., 2019). Finally, the present protocol includes only a passive control condition that precludes to extract and delineate the meditation-specific effects. Therefore, we cannot exclude that the main effects found in the meditation group could derive from an engagement in a repeated activity over a week or from expectancy for the meditation practice, rather than the merely practice of Focus Attention on the breath. An active control group, e.g. health enhancement programs (MacCoon et al., 2012), that can control variables such as social interaction with the group and teachers and amount of exercises, might have been beneficial in gaining insight into the specificity of the FA protocol applied here. In the future, it will be beneficial to include an active control condition for a proper interpretation of our results.

## CRediT authorship contribution statement

**Sara Trova**: Conceptualization, Methodology, Investigation, Formal analysis, Visualization, Data curation, Writing – original draft; **Yuki Tsuji**: Conceptualization, Methodology, Validation, Resources; **Haruka Horiuchi**: Investigation, Formal analysis; **Sotaro Shimada**: Conceptualization, Writing – review & editing, Supervision, Project administration.

## Data Statement

All data are available upon request to the first author of this manuscript.

## Acknowledgements

We would like to thank the Meiji University Institution (School of Science and Technology, Meiji University, 1-1-1 Higashi -Mita, Tama-ku, Kawasaki-shi, Kanagawa 214-8571, Japan) for providing a postdoctoral researcher fellowship allowing the current study to be carried out. Sincere thanks to Dr. Fabio Landuzzi (Postdoctoral Researcher at Aoyama Gakuin University, Department of Physics and Mathematics, Tokyo, Japan) for his contribution in the data analysis and for his kind support and encouragement, which help us in completion of this study.

